# A multi-modal machine learning approach towards predicting patient readmission

**DOI:** 10.1101/2020.11.20.391904

**Authors:** Somya D. Mohanty, Deborah Lekan, Thomas P. McCoy, Marjorie Jenkins, Prashanti Manda

## Abstract

Healthcare costs that can be attributed to unplanned readmissions are staggeringly high and negatively impact health and wellness of patients. In the United States, hospital systems and care providers have strong financial motivations to reduce readmissions in accordance with several government guidelines. One of the critical steps to reducing readmissions is to recognize the factors that lead to readmission and correspondingly identify at-risk patients based on these factors. The availability of large volumes of electronic health care records make it possible to develop and deploy automated machine learning models that can predict unplanned readmissions and pinpoint the most important factors of readmission risk. While hospital readmission is an undesirable outcome for any patient, it is more so for medically frail patients. Here, we develop and compare four machine learning models (Random Forest, XGBoost, CatBoost, and Logistic Regression) for predicting 30-day unplanned readmission for patients deemed frail (Age ≥ 50). Variables that indicate frailty, comorbidities, high risk medication use, demographic, hospital and insurance were incorporated in the models for prediction of unplanned 30-day readmission. Our findings indicate that CatBoost outperforms the other three models (AUC 0.80) and prior work in this area. We find that constructs of frailty, certain categories of high risk medications, and comorbidity are all strong predictors of readmission for elderly patients.

## I Introduction

The Healthcare Cost and Utilization Project (HCUP), estimates that unplanned 30-day readmission costs the United States $41.3 billion [1]. Approximately 18% of patients on Medicare were readmitted within 30 days of discharge - a number that remained relatively unchanged between 2007 and 2010 [2]. These unplanned hospital readmissions are both a burden on the US health care system as well as a strong indicator of sub-par quality of care [2].

Reducing unplanned readmissions is a goal that has significant health, wellness, and financial benefits for nations and their people. Given the strong health and financial implications, there has been a consistent push to understand and determine the key factors leading to these readmissions [2]. One of the first steps in reducing readmissions is to understand and determine the ever-evolving key causes that lead to instances of readmission, by developing predictive tools that assess risk of readmission. Consistent factors that lead to unplanned readmissions include premature discharge, length of stay in the hospital, type of hospital, lack of post-discharge treatments, and might include other factors [2].

While readmission within 30 days of discharge is an undesirable outcome for all patients, the outcomes can be particularly critical for the medically frail. Frailty, a syndrome which is marked by decreased physiological reserve, poor resilience, and increased vulnerability to stressors, is gaining recognition as an important risk factor and predictor of poor patient outcomes.

Another potential factor that might lead to unplanned readmission is the presence of one or more co-morbidities or the use of high risk medications. Comorbidity, defined as the coexistence of two or more medical conditions in a patient, is commonly included in risk prediction models to account for patient heterogeneity and chronic disease burden that is associated with more complex care and poor clinical outcomes [3].

The AGS Beers Criteria provides a list of medications that are either considered high risk or should be avoided by older adults in most circumstances [4]. These High Risk Medications (HRMs) continue to be prescribed and used in older patients which often leads to poor outcomes.

Many readmission risk prediction models exist [5, 6], but these models are limited in being able to predict readmission within a high risk population such as, acutely ill frail older adults with multiple comorbidities and medications. Our study adds to this literature and underscores the potential of Electronic Health Records (EHR) data as a rich clinical data source, while demonstrating the value of including frailty, high risk medications, and comorbidity in risk prediction models.

The goal of this study is to employ machine learning models that use a wide variety of data from electronic health records to predict 30-day unplanned readmission among patients deemed to be frail (aged 50 years or older). Specifically, we conducted a comparison of four parametric and non-parametric machine learning models to identify the best model for predicting readmission in older patients. We also investigated the merits of capturing various aspects of patient risk that are related to medical complexity and heterogeneity using novel indicators (frailty, comorbidity, and HRMs) extracted from EHR data on machine learning models. Finally we explore the most important factors that predict the risk of readmission.

### A. Background

Research indicates that frailty is associated with a range of adverse outcomes such as falls, functional and cognitive decline, disability, increased health care utilization, and premature mortality; thus, inclusion of frailty in risk prediction models is increasing. Estimates of the prevalence of frailty in the acute care setting ranges from 50% to 94% [7, 8].

Frailty can be measured in many ways but there are three well recognized methods. The Fried phenotype framework classifies frailty on the basis of the patient having at least three of five criteria (slow walking speed, weak grip strength, low physical activity, unintended weight loss, and exhaustion) [9]. The Rockwood deficit accumulation model identifies frailty based the number of deficits identified from the patient history and physical exams to calculate a frailty index (FI) score [10]. The FI consists of between 30 to 90 deficits (signs, symptoms, diseases, activities of daily living and disabilities, and physical and cognitive impairments). The third method is based on geriatric assessment and subjective determination of frailty by the health care provider and includes clinical tools such as the Clinical Frailty Scale (CFS), Identification of Seniors at Risk (ISR), and the Tilburg Frailty Indicator (TFI).

In the hospitalized older adult, the concept of frailty provides a framework for understanding vulnerabilities that render individuals more susceptible to the adverse effects of acute illness, inpatient treatments, and procedures during hospitalization that are independent of comorbidity [11]. Several recent studies utilize frailty in risk prediction models suggesting that frailty provides added value and offers a unifying approach to risk stratification for important outcomes in older adults [12, 13].

The presence of comorbidities can be represented using indices that account for specific subsets of diagnoses. A comorbidity index such as the Elixhauser Comorbidity Index (ECI) [14] aggregates selected medical diagnoses yielding a sum score. The Charlson Comorbidity Index (CCI), another widely used comorbidity index [15] contains 19 diseases including diabetes, congestive heart failure, etc. A more recent model of capturing comorbidities, Elixhauser Measure (ECI) [14] contains 31 conditions some of which are not accounted for in CCI [14].

Research on HRMs and hospital readmission is equivocal. In a study examining HRMs in hospitalized older adults, exposure to certain HRM classes such as benzodiazepines and opioids were associated with increased odds of readmission [16]. The American Geriatrics Society (AGS) Beers Criteria [17] for potentially inappropriate medication use in older adults is widely used by clinicians, healthcare workers, and researchers.

## II. Related Work

The use of machine learning for predicting readmission using patient demographic and medical records is well studied [2, 18, 19, 1, 20]. Some of these studies generated complex models that use thousands of features [2], while others limit themselves to patient information that can be easily collected within the hours of initial admission [18].

A comparison of commonly used models for predicting readmission risk studied a set of four models (LACE, STEPWISE logistic, LASSO logistic, and AdaBoost) [1]. The study finds that LACE has moderate predictive power with AUC scores around 0.65. Variables include, number of emergency room visits in the last year, Braden pressure ulcer risk score, polypharmacy, employment status, discharge disposition (patient’s anticipated location or status following a hospital visit), etc. LASSO was found to be the best model for both small and large data sizes (0.73 AUC).

Another study conducted on patients within the Maine Health Information Exchange created a risk model for identifying patients at risk for readmission within 30 days post discharge [2]. A risk assessment tool was developed on all patients in the system that clustered patients based on probability of readmission. The tool used 2000 features in addition to the number of days post discharge, to conduct a survival analysis using random forests. The model achieved a c-statistic (AUC) of 0.72 in predicting readmission.

While the above study uses a wide range of features and complex models to predict readmission, other studies have focused on identifying early hospital readmission factors in diverse patient populations using simple models with limited features [18]. In the study, data for 10,000 patients was grouped into four categories: sociodemographic factors, social support, health condition, and healthcare utilization. Logistic regression was used to determine significant predictors of 30-day readmission with an AUC of 0.61. Insurance status, marital status, having a regular physician, Charlson comorbidity index were among the best predictors for assessing risk of readmission.

Some studies augment EHR data available at the time of admission with administrative data available after patient discharge to build robust models that use diverse sources of information. In one such study, Tabak et al. [19], hypothesized that clinical severity and number of prior hospital stays increased a patient’s risk of readmission. The model was subsequently enhanced by using patient’s administrative data. A multi-variate Logistic Regression model was used to analyze the features and develop a risk model resulting in an AUC of 0.72.

The above studies predict readmission across all patient samples without discrimination for disease diagnoses leading to readmission. On the other hand, a number of studies have been conducted to identify risk factors for readmission after specific disease conditions. Goto et al. [21] use machine learning to predict 30-day readmission after hospitalization for Chronic Obstructive Pulmonary Disease (COPD). Patient characteristics and inpatient care data were used with Logistic Regression, Lasso Regression, and Deep Neural Network models to predict readmission after COPD. Logistic Regression was outperformed by both Lasso Regression and Deep Neural Networks; however all three models resulted in AUC of less than 0.62. Tube feeding duration, blood transfusion, use of thoracentesis, and sex were found to be important predictors for the machine learning models.

In another case of predicting readmission for specialized disease conditions, Mahajan et al. [20], explored Logistic Regression, Random Forest, Gradient Boost, and Neural Networks for predicting 30-day readmission for heart failure. Similarly, Awan et al [22] used machine learning to determine the most relevant predictors and variable transformations for predicting the risk of readmission or death following heart failure discharge. The team developed multi-layer perceptron models for predicting readmission using a set of 47 variables that included patient demographics, admission characteristics, medical history, socio-economic data, and medications (AUC = 0.66). A subset of eight variables was found to result similar predictive accuracy as compared to the full set of 47 variables. See [23] for research on predicting readmission for chronic or acute conditions like stroke, heart failure, myocardial infarction and pneumonia.

More recently, a study now considered the state of the art from scientists at Google, Stanford, and other institutions conducted an analysis of EHR to predict mortality, 30 day readmission, and prolonged length of stay [24]. The study reports that deep learning models were found to outperform traditional machine learning and clinical models at predicting the above events of interest. The models were validated using EHR data from two US academic medical centers and reported a 0.75-0.76 AUC for 30-day readmission.

## III. Methods

### A. Dataset, Data Cleaning, and Pre-processing

Data used in this analysis is from a collaborating health system in the Southeastern United States, and extracted from hospitals Epic® EHR patient data systems. Data is sourced across five hospitals each with a capacity of 85-535 beds. Data was collected for the time period of 2013-2017, and was filtered to only retain all admissions for adults 50 years and older with an inpatient stay lasting longer than 24 hours. The resulting dataset consists of 145,148 observations recorded from 76,294 patients admitted to the hospital. This data was de-identified in accordance with the data use agreement between UNC-Greensboro and the health system.

Figure 1 shows the steps of the data cleaning process. Starting with the raw dataset of 145,148 observations (76,294 patients), the first step is to remove any observations where the patients age was lower than 50, resulting in a removal of 1,327 observations. In order to create a dataset for 30 day readmission, in the second step we created a buffer of 30 days at the start and end of earliest and latest admission times in the data. Third, fourth, and fifth steps removed observations which had, less than 24 hours for length of stay in the hospital, died on initial admission, and had planned readmissions respectively. Sixth step removed any subsequent observations where the patient expired. Finally, we removed any patients where the disease diagnosis was related to pregnancy, childbirth, and postpartum care. The final dataset had 124,480 observations recorded from 68,152 patients. Within the dataset, 18,014 observations had subsequent unplanned readmissions, and 106,466 did not.

**Fig. 1:**
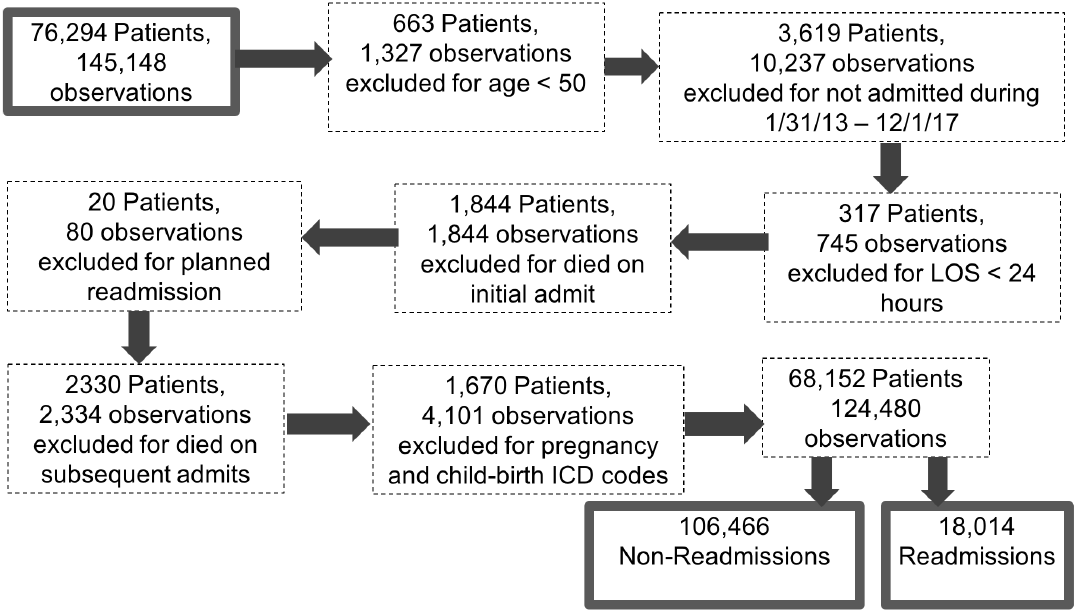
Steps and Exclusion criteria employed to prepare the EHR data for modeling

### B. Variable Extraction

The dataset contains 81 variables (Table I) that can be divided into six categories: 1) Frailty, 2) Comorbidity, 3) High Risk Medications, 4) Disease Diagnosis, 5) Demographic, 6) Healthcare and Insurance Utilization.

**TABLE I:** List of variables and their corresponding category utilized in predicting 30-day readmission risk.

| Category | Variable |
| --- | --- |
| Frailty | FRS-26-ICD; Malnutrition; Abnormal Weight; Dysphagia; Delirium; Dementia; Depression; Vision; Weakness; Fatigue; Dyspnea; Difficulty Walking; Falls; Chronic Pain; Urine incontinence; Fecal incontinence; Decubitus Ulcer; PImaterial Material Resources; Social Support Problems; Smoking; WBC; Albumin; C-reactive protein; Hemoglobin; Glucose; Creatinine; Sodium |
| Comorbidity | Elixhauser Comorbidity Index |
| High Risk Medications | Anticholinergics, Antispasmodics; Benzodiazepines Nonbenzodiazepine Hypnotics; Cardiovascular; Central Nervous System; Endocrine; Pain Meds; H2 Receptor Blockers, Proton Pump Inhibitors; Antipsychotics; Anti-infective; Genitourinary |
| Disease Diagnosis (ICD10) | Infectious and Parasitic Diseases; Neoplasms; Disease of the blood and blood-forming organs and certain disorders involving the Endocrine, Nutritional, and Metabolic Diseases; Mental, Behavioral and Neurodevelopmental disorders; Diseases of the Nervous System; Diseases of the Eye and Adnexa; Diseases of the Ear and Mastoid Process; Diseases of the Circulatory System; Diseases of the Respiratory System; Diseases of the Digestive System; Diseases of the Skin and Subcutaneous Tissue; Diseases of the Musculoskeletal System and Connective Tissue; Diseases of Genitourinary System; Pregnancy, Childbirth, and the Puerperium; Certain Conditions Originating in the Perinatal Period; Congenital malformations, deformations, and chromosomal abnormalities; Symptoms, signs, and abnormal clinical and laboratory findings, not elsewhere classified; Injury, poisoning, and certain other consequences of external causes; External Causes of Morbidity; Factors influencing health status and contact with health services; Num Diagnosis; Num Unique Diagnosis |
| Demographic | Age; Sex; Race/ethnicity; Marital Status |
| Health Care and Insurance Utilization | Num Prior Readmits; Quad Length of Stay; Hospital Service; Admit Source; Discharge Disposition; Primary Payor; Secondary Payor; Hospital (deidentified, 1-5); Hospital Unit; Hospital Admit Time |

#### Frailty

Our study utilized a proxy measure for frailty (FRS-26-ICD) drawn from International Classification of Disease, Tenth Revision, Clinical Modification (ICD-10-CM) disease diagnosis codes that encompass, common geriatric syndromes, psycho-social factors, and blood biomarkers. The FRS-26-ICD defines frailty as a clinical syndrome resulting from multi-system physiologic impairments and failed integrative responses with diminished capacity to resist and recover from stressors [25, 26]. In addition to the FRS-26-ICD composite score, 32 additional variables were incorporated to capture the measure of frailty for every patient. These variables include but are not limited to Malnutrition, Abnormal Weight, Fatigue, Difficulty Walking, etc. Laboratory values were discretized based on a reference range that indicates risk. Original indicators of frailty compiled from blood biomarkers were represented using laboratory reference ranges for abnormal (high, or low) for factors such as albumin, hemoglobin, sodium, and white blood cells.

#### Comorbidity Indices

Both Charlson Comorbidity (CCI) and Elixhauser comorbidity indices were extracted and calculated for each observation using the ICD diagnosis codes present within the data.

#### High Risk Medications

Using the AGS Beers Criteria and the prescribed drugs extracted from the EHR data, each observation was marked for its use of High Risk Medication (HRM). Specifically, 10 different categories were developed based on type of medication and/or the organ system targeted by the HRM in the human body. Each observation was marked containing the HRM category if the prescribed drugs matched the criteria. Specific drug names were mapped to generic names, and a string matching algorithm was used to link the drugs to the HRMs.

#### Disease Diagnosis

ICD-10 codes [27] are recorded in the EHR data to describe the principal problem attributed to the hospital admission as well as any additional diagnoses for each patient not directly related to the principal problem. Each ICD-10 code was grouped into 19 high level classifications as described by the CDC guidelines [27]. For each observation, the principal and additional diagnosis were mapped to the high level classification. Observations which did not contain disease diagnosis were marked ‘NA’ (Not Available). The mapping of ICD-10 codes to the corresponding higher level category was performed to reduce the dimensionality for machine learning.

#### Demographic

Several patient level variables recorded at the time of admission were extracted from the EHR data. These include, Age, Sex, Ethnic Group, Patient Race, and Marital Status. Age was further discretized into 20 groups using frequency histograms.

#### Healthcare and Insurance Utilization

Use of hospital Service, hospital, hospital unit, admit source, discharge disposition, along with insurance payor information were extracted from the administrative records. Additionally, the admission time was discretized into 4 groups 6 hours apart: Morning, Afternoon, Evening, and Late night. For each observation, number of prior admits and readmissions were calculated.

#### Class Label

For each observation, a time delta 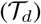 was calculated by taking the difference between the discharge time to the next admit time for the same patient. If the 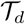 was within 30 days, the observation was marked as *Class_Label* = 1 (readmission), else *Class_Label* = 0 (non-readmission) if the time delta 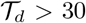 days. Pairwise collinearity tests were conducted for all pairs of features in the data to eliminate highly correlated features. Following this, one-hot encoding was performed on the categorical features resulting in the dataset expanding to 369 dimensions.

### C. Machine Learning Models

After data cleaning and variable extraction, the resulting dataset consists of 122,813 observations with 369 variables. Out of this 18,014 were readmissions *(Class_Label =* 1) and 106,466 non-readmissions *(Class_Label =* 0). This data was split into a 80:20 ratio, with 80% of the data being used for development of models, while the rest 20% was used for validation of the models. Three different strategies were used to address the imbalance between the two target classes *(Class_Label =<* 1,0 >): under sampling, over sampling, and no sampling. Under sampling selects a same sized subsample from the majority class *(Class_Label =* 0) with respect to the size of the minority class *(Class_Label =* 1). The resulting dataset has a balanced number of observations between the two classes.

Over sampling was done with Synthetic Minority Oversampling TEchnique (SMOTE) [28], which oversamples on minority instances by synthesizing new data points between real data instances. SMOTE oversampling was performed only on the training data.

Four models were developed and compared: 1) Random Forest, 2) XGBoost, 3) Category Boost, and 4) Logistic Regression. **Random Forest** Random Forest is an ensemble prediction method that consists of a set of individual decision trees [29]. The decision trees are designed to have low correlation to each other to encourage diversity among the trees. The prediction of individual trees is aggregated to determine the prediction of the random forest. Random Forests use the principles of Bootstrapping and Aggregating to build trees based on different subsets of the training data using different subsets of features.

**XGBoost** XGBoost is an implementation of gradient boosted decision trees whose main advantages are execution speed and model performance [30]. Boosted decision trees use boosting - an ensemble technique where each tree or model corrects errors made by previous trees. The technique adds more and more trees/models until the overall accuracy cannot be improved anymore. The predictions of the models are added together to make the final prediction.

**CatBoost** CatBoost stands for Category Boosting [31] - a variation of the boosting techniques typically used for machine learning. CatBoost improves on existing boosting techniques by using ordered boosting and a novel algorithm for effectively processing categorical features. Empirical studies have shown that CatBoost outperforms other publicly available boosting algorithms. CatBoost does not require any special pre-processing for categorical features such as encodings etc. Instead, the algorithm converts categorical values into numbers using statistics on combinations of categorical features. The algorithm has been shown to be more robust therefore reducing the need for parameter tuning and optimization. It also reduces the chance of overfitting.

**Logistic** Regression Logistic regression is one of the most widely used models for analyzing EHRs. An extension of linear regression, a basic logistic regression determines the probability of classification problems with a binary outcome [32]. The model uses a logistic function to fit a linear equation between 0 and 1. Each of models were evaluated using k-fold (k = 10) cross-validation on Precision, Recall, F-1, and AUC, for their performance metrics.

## IV. Results and Discussion

The EHR dataset prior to pre-processing and manipulation contained 145,148 observations corresponding to 76,294 patients. A series of seven exclusion criteria (Figure 1) were sequentially applied to result in 124,480 observations for 68,152 patients. These observations consisted of 18,014 readmissions and 106,466 non-readmissions.

The four machine learning models were applied to data created using under sampling after exploring no sampling, over sampling strategies. Of the four models tested (Figure 2), Category Boosting outperformed the other models with an AUC of 0.80. Random Forest and XGBoost trailed behind Category Boosting with AUCs of 0.78 and 0.77 respectively. Logistic Regression, which is widely used for prediction of readmission, performed substantially worse as compared to the other three models with an AUC of 0.69.

**Fig. 2:**
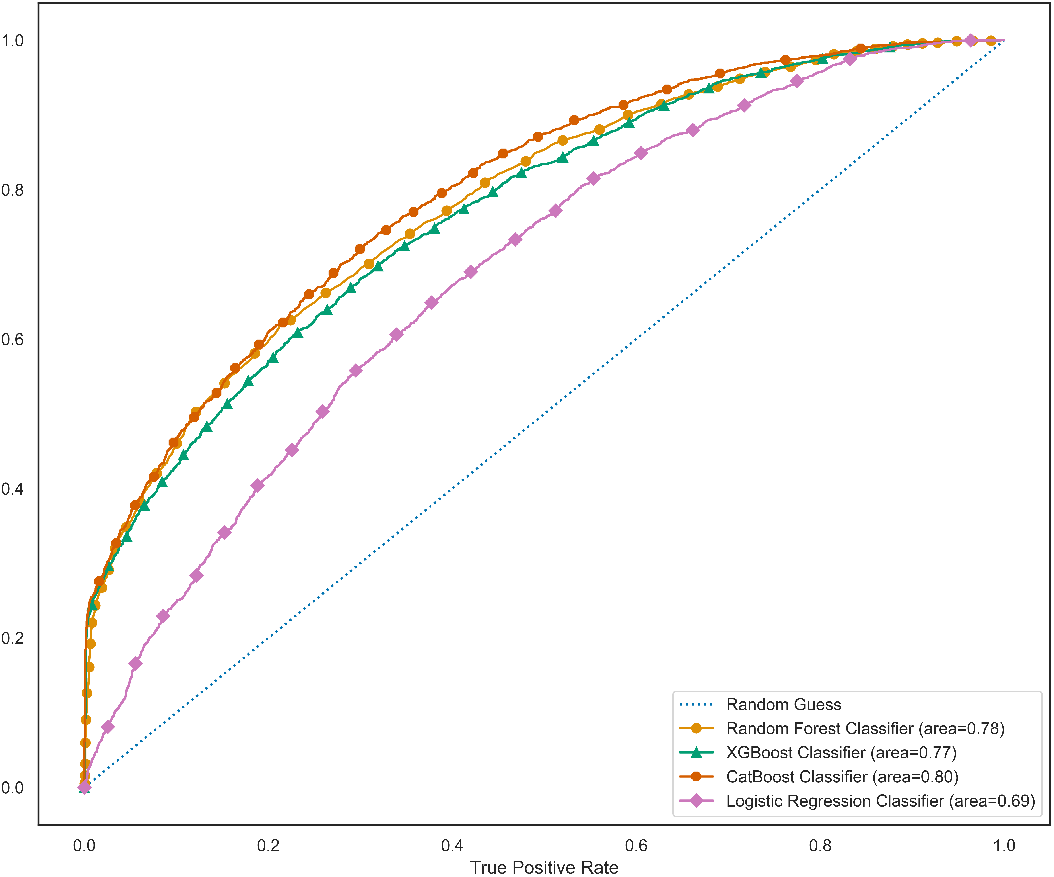
Comparison of four models for prediction of unplanned 30-day readmission with data created using under sampling.

**Fig. 3:**
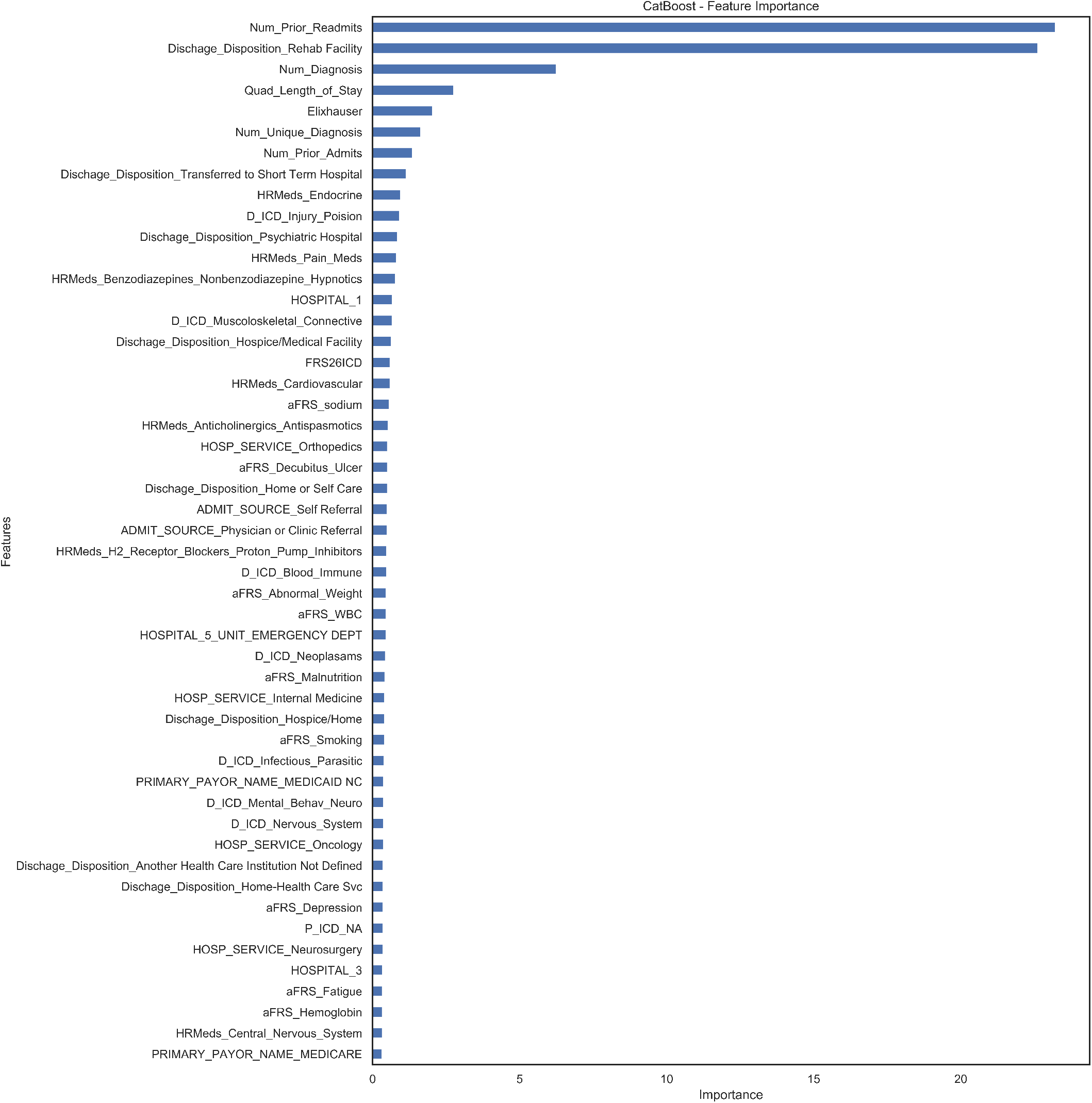
Important features for prediction readmission by CatBoost model.

We further explored the best model from our tests (Category Boosting) to determine the most important features for predicting 30-day readmission. The most important factor was found to be the number of prior readmissions followed by discharge from a rehab facility. Morbidity and comorbidity, represented by the number of diagnoses, number of unique diagnoses, and ECI were also found to be an important factor. Next, we see that the length of stay and Elixhauser comorbidity index impact the risk of readmission. These findings are in line with a systematic review [6] that found that among 60 studies, comorbidities, LOS, and number of previous admissions were the most frequently cited predictors for readmission. Discharge disposition was also a strong feature, including discharge to a short-term hospital and psychiatric hospital (compared to discharge to home or self care which was not a strong feature of readmission). Research also indicates that the type of index admission such as emergency department admissions (here referred to Admit Source-Self Referral) are strongly associated with readmission [3]

An unexpected finding was that almost all patients who were discharged to a rehabilitation hospital were readmitted within 30 days (844 of 925 patients, 92%); this was among the strongest features associated with readmission. This clinical setting accommodates patient with greater medical complexity requiring professional nursing and rehabilitation services for problems such as hip fracture and stroke recovery. Complex care transitions and greater risk for clinical instability after acute care discharge in this patient population may contribute to early hospital readmission [33]. Further investigation of patients discharged to a rehabilitation hospital is warranted to identify contributing factors and strategies to improve discharge planning and care transitions.

Findings from the ML models in our study confirm the importance of comorbidity in readmission prediction, but also highlights the relevance of both comorbidity and frailty in readmission. We found, there was no collinearity between Frailty and comorbidity (ECI) and frailty (FRS-26-ICD) as measured by Pearson correlations.

In our study, frailty as measured by the FRS-26-ICD was a strong feature in the ML model for 30-day readmission. These findings contrast with results from several systematic reviews that highlight the uneven quality and low predictive accuracy of frailty instruments applied in prediction models in the acute care setting [34]. The FRS-26-ICD used in the present study is at risk for for under-coding since coding for ambiguous syndromes such as weakness, fatigue, and dysphagia, which are indicators in the FRS-26-ICD, may not be adequately recorded by the healthcare provider.

Diagnosis coding fields are limited in number, further restricting codes that may more comprehensively and timely characterize patient health status. Individual FRS indicators were also strong features in the ML model, and were included in the analyses based on the rationale that no single indicator has a major influence on the FRS score, and that the composite measure is greater than the sum of its parts. The presence of 9 individual FRS indicators in the top features (e.g., abnormal weight, smoking, sodium, fatigue, malnutrition corresponding to the well-known frailty phenotype defined by [35]), indicates the importance of including indicators of frailty for prediction of readmission. Incorporating frailty in risk prediction can help account for these vulnerabilities and help identify the highest risk population in need of tailored interventions for inpatient care, discharge planning and care transitions to prevent readmission and other adverse health outcomes. The inclusion of frailty in predictive modeling for 30-day all cause readmission is a key contribution of this work and our results highlight the implications of using frailty indicators to reduce the risk of readmission. These results also underscore the importance of deepening our understanding and study of frailty.

One systematic review [6] found that among 60 studies with 73 unique models for 28-day or 30-day unplanned readmission, comorbidities, LOS, and number of previous admissions were the most frequently cited predictors. Alternatively, laboratory tests and medications were more frequently included in the models for disease/medical conditions, mental health, and surgical conditions related unplanned readmissions. The review by Zhou et al [6] also suggests that compared to variables that are extracted from claims data (e.g., comorbidities, LOS, number of previous admissions), the addition of clinical data such as laboratory tests and medications were also valuable in prediction models. Our study which included frailty, blood biomarkers and HRMs, contributes new information to the literature for all-cause 30-day readmission.

Seven high risk medication categories were found to be important features for predicting readmission. These include Endocrine, Pain Medications, Cardiovascular, Anto-cholinergics/Antispasmodics, Receptor blockers/Proton pump inhibitors, Benzodiazepines/Hypnotics, and Central Nervous System. The strong presence of these HRM categories in the feature importance may reflect illness severity and multimorbidity in the acute care population and may provide added value in estimates of readmission risk. In the acutely ill, this is important since the presence of multiple chronic conditions and other interventions can lead to a “prescribing cascade” in which a medication prescribed for a medical problem causes adverse effects that are misinterpreted as signs and symptoms of another medical problem [36]. Many factors are associated with the risk of readmission, such as, age, race/ethnicity, comorbidity and disease severity, health care utilization and length of stay, are non-modifiable attributes of a patient. Appropriate pharmacotherapy in older adults, especially in those with frailty and multimorbidity, requires a balance between risks and benefits of medications.

### A. Limitations

The retrospective nature of our study limited analyses to the information available in the EHR and is dependent upon accurate and timely documentation by providers and coders. Data coding typically occurs after hospitalization (for principle diagnosis or reason for admission, LOS) thus, the utility of this measure is limited as a real-time clinical decision support tool. As hospital health informatics evolve, generation of a frailty risk score could be incorporated into future EHRs and made available in real time to clinicians caring for patients based on their prior hospitalizations. However, using historical data and modeling patients with similar clinical and health care utilization patterns, can prove to be useful as an admission screen tool. This can readily identify patients fitting those risk groups, and more timely proactive measures can be initiated during hospitalization. The 30-day readmission rate in this study may be lower than the actual admission rate because the scope of our study did not have readmission data from hospitals outside of the health system. Our analyses only included medications identified as high risk according to Beers Criteria and taken during the hospitalization, which does not account for other medications taken before hospitalization that may also exert adverse effects. Our data is also based on healthcare data and coding practices in a single health system in the U.S. and may not be applicable in other settings. Despite these limitations, this study offers a prediction model for 30-day readmission based on a comprehensive set of variables from EHR. Future research is focused on using clinical flowsheet data and unstructured text derived from nursing and other health professional documentation within the EHR.

## V. Conclusions

This paper reports results for a machine learning based risk prediction model using EHR data for 30-day unplanned hospital readmission with good predictability that included sociodemographic data and comorbidity and less commonly used variables such as frailty and HRMs. We find that, in a comparison of four models, Category boosting substantially outperforms the often used Logistic Regression at the task of predicting readmission. Using readily available EHR data is an efficient way to identify high and low risk patients and focus transitional care programs and strategies to help reduce early hospital readmissions. Factors such as frailty, high risk medication, and multi-morbidity are strong features in our readmission ML models and can guide individual tailored multidisciplinary care and discharge planning to prevent readmissions.

## References

[1] L. Tong, C. Erdmann, M. Daldalian, J. Li, and T. Esposito, “Comparison of predictive modeling approaches for 30-day all-cause non-elective readmission risk,” BMC medical research methodology, vol. 16, no. 1, p. 26, 2016.

[2] S. Hao, Y. Wang, B. Jin, A. Y. Shin, C. Zhu, M. Huang, L. Zheng, J. Luo, Z. Hu, C. Fu et al., “Development, validation and deployment of a real time 30 day hospital readmission risk assessment tool in the maine healthcare information exchange,” PloS one, vol. 10, no. 10, 2015.

[3] P. Zhao and I. Yoo, “A systematic review of highly generalizable risk factors for unplanned 30-day all-cause hospital readmissions,” J. Heal. Med. Informatics, vol. 8, no. 04, 2017.

[4] D. Fick, T. Semla, M. Steinman, J. Beizer, N. Brandt, and R. Dombrowski, “American geriatrics society beers criteria® update expert panel, et al. american geriatrics society 2019 updated ags beers criteria® for potentially inappropriate medication use in older adults,” J Am Geriatr Soc, vol. 67, no. 4, pp. 674–94, 2019.

[5] D. Kansagara, H. Englander, A. Salanitro, D. Kagen, C. Theobald, M. Freeman, and S. Kripalani, “Risk prediction models for hospital readmission: a systematic review,” Jama, vol. 306, no. 15, pp. 1688–1698, 2011.

[6] H. Zhou, P. R. Della, P. Roberts, L. Goh, and S. S. Dhaliwal, “Utility of models to predict 28-day or 30-day unplanned hospital readmissions: an updated systematic review,” BMJ open, vol. 6, no. 6, p. e011060, 2016.

[7] P. Forti, F. Maioli, E. Zagni, T. Lucassenn, L. Montanari, B. Maltoni, G. L. Pirazzoli, G. Bianchi, and M. Zoli, “The physical phenotype of frailty for risk stratification of older medical inpatients,” The journal of nutrition, health & aging, vol. 18, no. 10, pp. 912–918, 2014.

[8] D. Basic and C. Shanley, “Frailty in an older inpatient population: using the clinical frailty scale to predict patient outcomes,” Journal of aging and health, vol. 27, no. 4, pp. 670–685, 2015.

[9] L. P. Fried, L. Ferrucci, J. Darer, J. D. Williamson, and G. Anderson, “Untangling the concepts of disability, frailty, and comorbidity: implications for improved targeting and care,” The Journals of Gerontology Series A: Biological Sciences and Medical Sciences, vol. 59, no. 3, pp. M255–M263, 2004.

[10] S. D. Searle, A. Mitnitski, E. A. Gahbauer, T. M. Gill, and K. Rockwood, “A standard procedure for creating a frailty index,” BMC geriatrics, vol. 8, no. 1, p. 24, 2008.

[11] B. Santos-Eggimann, P. Cuénoud, J. Spagnoli, and J. Junod, “Prevalence of frailty in middle-aged and older community-dwelling europeans living in 10 countries,” The Journals of Gerontology: Series A, vol. 64, no. 6, pp. 675–681, 2009.

[12] D. H. Kim, E. Patorno, A. Pawar, H. Lee, S. Schneeweiss, and R. J. Glynn, “Measuring frailty in administrative claims data: Comparative performance of four claims-based frailty measures in the us medicare data,” The Journals of Gerontology: Series A, vol. 75, no. 6, pp. 1120–1125, 2020.

[13] J. B. Segal, H.-Y. Chang, Y. Du, J. Walston, M. Carlson, and R. Varadhan, “Development of a claims-based frailty indicator anchored to a well-established frailty phenotype,” Medical care, vol. 55, no. 7, p. 716, 2017.

[14] A. Elixhauser, C. Steiner, D. R. Harris, and R. M. Coffey, “Comorbidity measures for use with administrative data,” Medical care, pp. 8–27, 1998.

[15] M. E. Charlson, P. Pompei, K. L. Ales, and C. R. MacKenzie, “A new method of classifying prognostic comorbidity in longitudinal studies: development and validation,” Journal of chronic diseases, vol. 40, no. 5, pp. 373–383, 1987.

[16] J. M. Pavon, Y. Zhao, E. McConnell, and S. N. Hastings, “Identifying risk of readmission in hospitalized elderly adults through inpatient medication exposure,” Journal of the American Geriatrics Society, vol. 62, no. 6, pp. 1116–1121, 2014.

[17] A. G. S.. B. C. U. E. Panel, D. M. Fick, T. P. Semla, J. Beizer, N. Brandt, R. Dombrowski, C. E. DuBeau, W. Eisenberg, J. J. Epplin, N. Flanagan et al., “American geriatrics society 2015 updated beers criteria for potentially inappropriate medication use in older adults,” Journal of the American Geriatrics Society, vol. 63, no. 11, pp. 2227–2246, 2015.

[18] O. Hasan, D. O. Meltzer, S. A. Shaykevich, C. M. Bell, P. J. Kaboli, A. D. Auerbach, T. B. Wetterneck, V. M. Arora, J. Zhang, and J. L. Schnipper, “Hospital readmission in general medicine patients: a prediction model,” Journal of general internal medicine, vol. 25, no. 3, pp. 211–219, 2010.

[19] Y. P. Tabak, X. Sun, C. M. Nunez, V. Gupta, and R. S. Johannes, “Predicting readmission at early hospitalization using electronic clinical data: an early readmission risk score,” Medical care, vol. 55, no. 3, p. 267, 2017.

[20] S. Mahajan and R. Ghani, “Using ensemble machine learning methods for predicting risk of readmission for heart failure.” Studies in health technology and informatics, vol. 264, pp. 243–247, 2019.

[21] T. Goto, T. Jo, H. Matsui, K. Fushimi, H. Hayashi, and H. Yasunaga, “Machine learning-based prediction models for 30-day readmission after hospitalization for chronic obstructive pulmonary disease,” COPD: Journal of Chronic Obstructive Pulmonary Disease, vol. 16, no. 5-6, pp. 338–343, 2019.

[22] S. E. Awan, M. Bennamoun, F. Sohel, F. M. Sanfilippo, B. J. Chow, and G. Dwivedi, “Feature selection and transformation by machine learning reduce variable numbers and improve prediction for heart failure readmission or death,” PloS one, vol. 14, no. 6, 2019.

[23] C. Eckert, N. Nieves-Robbins, E. Spieker, T. Louwers, D. Hazel, J. Marquardt, K. Solveson, A. Zahid, M. Ahmad, R. Barnhill et al., “Development and prospective validation of a machine learning-based risk of readmission model in a large military hospital,” Applied clinical informatics, vol. 10, no. 02, pp. 316–325, 2019.

[24] A. Rajkomar, E. Oren, K. Chen, A. M. Dai, N. Hajaj, M. Hardt, P. J. Liu, X. Liu, J. Marcus, M. Sun et al., “Scalable and accurate deep learning with electronic health records,” NPJ Digital Medicine, vol. 1, no. 1, p. 18, 2018.

[25] D. A. Lekan, D. C. Wallace, T. P. McCoy, J. Hu, S. G. Silva, and H. E. Whitson, “Frailty assessment in hospitalized older adults using the electronic health record,” Biological research for nursing, vol. 19, no. 2, pp. 213–228, 2017.

[26] L. Rodríguez-Mañas and A. Sinclair, “Frailty: The quest for new domains, clinical definitions and subtypes. is this justified on new evidence emerging?” The journal of nutrition, health & aging, vol. 18, no. 1, p. 92, 2014.

[27] C. for Medicare & Medicaid Services et al., “Icd-10-cm official guidelines for coding and reporting,” 2012.

[28] N. V. Chawla, K. W. Bowyer, L. O. Hall, and W. P. Kegelmeyer, “Smote: synthetic minority over-sampling technique,” Journal of artificial intelligence research, vol. 16, pp. 321–357, 2002.

[29] T. Shi and S. Horvath, “Unsupervised learning with random forest predictors,” Journal of Computational and Graphical Statistics, vol. 15, no. 1, pp. 118–138, 2006.

[30] T. Chen, T. He, M. Benesty, V. Khotilovich, and Y. Tang, “Xgboost: extreme gradient boosting,” R package version 0.4-2, pp. 1–4, 2015.

[31] L. Prokhorenkova, G. Gusev, A. Vorobev, A. V. Dorogush, and A. Gulin, “Catboost: unbiased boosting with categorical features,” in Advances in neural information processing systems, 2018, pp. 6638–6648.

[32] D. G. Kleinbaum, K. Dietz, M. Gail, M. Klein, and M. Klein, Logistic regression. Springer, 2002.

[33] S. L. Shih, P. Gerrard, R. Goldstein, J. Mix, C. M. Ryan, P. Niewczyk, L. Kazis, J. Hefner, D. C. Ackerly, R. Zafonte et al., “Functional status outperforms comorbidities in predicting acute care readmissions in medically complex patients,” Journal of general internal medicine, vol. 30, no. 11, pp. 1688–1695, 2015.

[34] O. Theou, E. Squires, K. Mallery, J. S. Lee, S. Fay, J. Goldstein, J. J. Armstrong, and K. Rockwood, “What do we know about frailty in the acute care setting? a scoping review,” BMC geriatrics, vol. 18, no. 1, p. 139, 2018.

[35] L. P. Fried, C. M. Tangen, J. Walston, A. B. Newman, C. Hirsch, J. Gottdiener, T. Seeman, R. Tracy, W. J. Kop, G. Burke et al., “Frailty in older adults: evidence for a phenotype,” The Journals of Gerontology Series A: Biological Sciences and Medical Sciences, vol. 56, no. 3, pp. M146–M157, 2001.

[36] P. A. Rochon and J. H. Gurwitz, “Drug therapy,” The Lancet, vol. 346, no. 8966, pp. 32–36, 1995.

